# Dengue virus exploits the host tRNA epitranscriptome to promote viral replication

**DOI:** 10.1101/2023.11.05.565734

**Authors:** Cheryl Chan, Newman Siu Kwan Sze, Yuka Suzuki, Takayuki Ohira, Tsutomu Suzuki, Thomas J. Begley, Peter C. Dedon

## Abstract

The 40-50 RNA modifications of the epitranscriptome regulate posttranscriptional gene expression. Here we show that flaviviruses hijack the host tRNA epitranscriptome to promote expression of pro-viral proteins, with tRNA-modifying ALKBH1 acting as a host restriction factor in dengue virus infection. Early in the infection of human Huh-7 cells, ALKBH1 and its tRNA products 5-formylcytidine (f^5^C) and 2’-*O*-methyl-5-formylcytidine (f^5^Cm) were reduced. ALKBH1 knockdown mimicked viral infection, but caused increased viral NS3 protein levels during infection, while ALKBH1 overexpression reduced NS3 levels and viral replication, and increased f^5^C and f^5^Cm. Viral NS5, but not host FTSJ1, increased f^5^Cm levels late in infection. Consistent with reports of impaired decoding of leucine UUA codon by f^5^Cm-modified tRNA^Leu(CAA)^, ALKBH1 knockdown induced translation of UUA-deficient transcripts, most having pro-viral functions. Our findings support a dynamic ALKBH1/f^5^Cm axis during dengue infection, with virally-induced remodeling of the proteome by tRNA reprogramming and codon-biased translation.

---

Human cells possess innate antiviral defense mechanisms such as the interferon response system and restriction factors that detect and limit viral replication during infection [1–4]. This is exemplified by well-characterized restriction factors for human immunodeficiency virus 1 (HIV-1), including TRIM5α, TRIMCyp, and members of the APOBEC family of mRNA cytidine deaminases [5–7]. At the same time, as obligate intracellular entities, viruses have evolved a variety of strategies to evade the host defense and to hijack host factors to successfully propagate [8–10]. Dengue viruses (DENV) are positive-stranded single-strand RNA viruses of the family Flaviviridae and represent some of the most common and clinically important arboviral human pathogens for which no drugs or effective vaccines are available [11, 12].

Examples of host factors involved in dengue infection include attachment molecules C-type lectin DC-SIGN and glycosaminoglycans, intracellular cholesterol transporters, and the replication factor DDX3 that influence DENV infectivity and replication [13–17]. Identification and validation of host cellular factors that influence virus infection are essential to expanding the target space for developing host-directed antiviral (HDA) therapies [10, 18, 19].

The epitranscriptome – the system of post-transcriptional modifications of all forms of RNA in an organism – is emerging as an important layer of regulatory control at the interface of virus-host interactions and its dysregulation has been linked to many human diseases [20]. This is illustrated by *N^6^*-methyladenosine (m^6^A), which is installed by cellular “writer” METTL3-METTL14 methyltransferases, removed by “eraser” FTO and ALKBH5 demethylases, and recognized by a variety of m^6^A “readers” that regulate the fate of the target RNA. m^6^A is present in both cellular and viral RNA and has functions in the virus life cycle as well as the cell’s response to viral infection [21–23]. In *Flaviviridae* infection, for example, m^6^A negatively regulates Zika virus replication by directly modulating the binding of m^6^A reader YTHDF proteins to viral RNA as well as indirectly by altering the m^6^A landscape of cellular mRNAs [24]. The m^6^A modification also suppresses hepatitis C virus (HCV) infectious particle production (but not HCV translation and RNA replication) largely by modulating binding of YTHDF proteins to m^6^A-containing HCV RNA at sites of HCV particle production [25].

More broadly [26], aberrations in the activity of highly conserved writer, reader, and eraser enzymes have been linked to disease [27]. For example, in human urothelial carcinoma of the bladder, overexpression of the methyltransferase NSUN2 leads to high levels of 5-methylcytosine (m^5^C) in oncogene messenger RNAs (mRNA), which are then stabilized by binding of the m^5^C reader, YBX1 [28]. Parallel observations in transfer RNAs (tRNA) further illustrate the role of aberrant modifications in promoting disease. Elevated levels of the human tRNA writers ELP3/CTU1/CTU2 that catalyze modification of tRNA wobble uridines have been shown to support melanoma cell survival and drive resistance to MAPK therapeutic agents through a translation-dependent mechanism [29]. Another well-developed model for cancer-driving tRNA modifications and codon-biased translation involves over-expression of METTL1 in acute myelogenous leukemia, gliomas, sarcomas, and other cancers [30–32]. The m^7^G46 modification catalyzed by METTL1 stabilizes tRNAs whose codons are enriched in mRNAs for cancer-driving cell proliferation genes [32].

One mechanism linking defects in the tRNA epitranscriptome to translational dysfunction and pathobiology involves altered translational efficiency of families of transcripts possessing biased usage of codons matching the modification-altered tRNAs [33–39]. This mechanism has been observed in a wide range of organisms. For example, mycobacteria respond to hypoxia by increasing uridine 5-oxyacetic acid (cmo^5^U) at the wobble position of tRNA^Thr(UGU)^ to more efficiently translate stress response transcripts enriched with ACG codons, while mRNAs enriched with the synonymous ACC codon, the so-called optimal codon for Thr, have reduced translational efficiency [36]. In response to oxidative stress, yeast cells rely on the wobble uridine tRNA writer, Elongator, to optimize translation of highly-expressed transcripts enriched with AAA and essential for oxidative stress survival [39], while yeast respond to alkylation stress by increasing Trm9-mediated formation of mcm^5^U on tRNA^Arg(UCU)^ that drives translation of mRNAs enriched in the cognate codon AGA [33]. In mice, tRNA methyltransferase ALKBH8 is required for the formation of 5-methoxycarbonylmethyl-2’-O-methyluridine (mcm^5^Um)-dependent translation of oxidative stress-response selenoproteins [38]. Finally, over expression of the m^7^G46 writer METTL1 in several human cancers leads to increased levels of one of its substrate tRNAs, tRNA-Arg-TCT-4-1, and increased translation of oncogenic mRNAs enriched with the cognate codon AGA [32].

Given the significantly different codon usage patterns in viral and human genomes [26], the observations and models for codon-biased translation driven by tRNA modifications raises the question of epitranscriptomic regulation of viral replication during infections. More specifically, the stress-regulated translation model predicts that host cells will respond to the stress of viral infections by tRNA modification-mediated translation of stress-response genes, while the virus could co-opt the host translational machinery to selectively translate its own proteins that arise from genes with strikingly different codon usage compared to humans [26]. Here we report that the multifunctional DNA and RNA writer and eraser, ALKBH1, behaves as a host restriction factor that is suppressed during dengue virus infection, while virally-encoded NS5 protein modifies ALKBH1 target tRNAs to facilitate translation of codon-biased pro-viral transcripts. The results point to the epitranscriptome as a potentially rich source of targets for developing antiviral therapeutics.

## Results

### A model system for epitranscriptome-regulated DENV infection

To investigate the role of the tRNA epitranscriptome during DENV infection, we first established an efficient (>50% infection) system for DENV serotype 2 new guinea C strain (DENV2 NGC) infection of human liver Huh-7 cells in culture (**Fig. 1a**). Fluorescence-activated cell sorting (FACS) analysis at 24 h post-infection (24 hpi) using an antibody against dengue E protein 4G2 showed 50-70% of Huh-7 cells were consistently infected at a low multiplicity of infection (MOI 1), with viral NS3 protein maximally expressed at 24 hpi (**Fig. 1a**). This infection system therefore afforded high virus infectivity of cells to minimize signal dilution by uninfected cells, minimized the potential for stress artefacts caused by cell sorting to enrich for infected cells, and represented a physiologically relevant cell type for human DENV infection. Paired mock-and DENV2-infected cells were then monitored for morphological changes and cell death during the course of infection, until cell death was evident at 48 hpi (**Fig. S1**).

**Figure 1.**
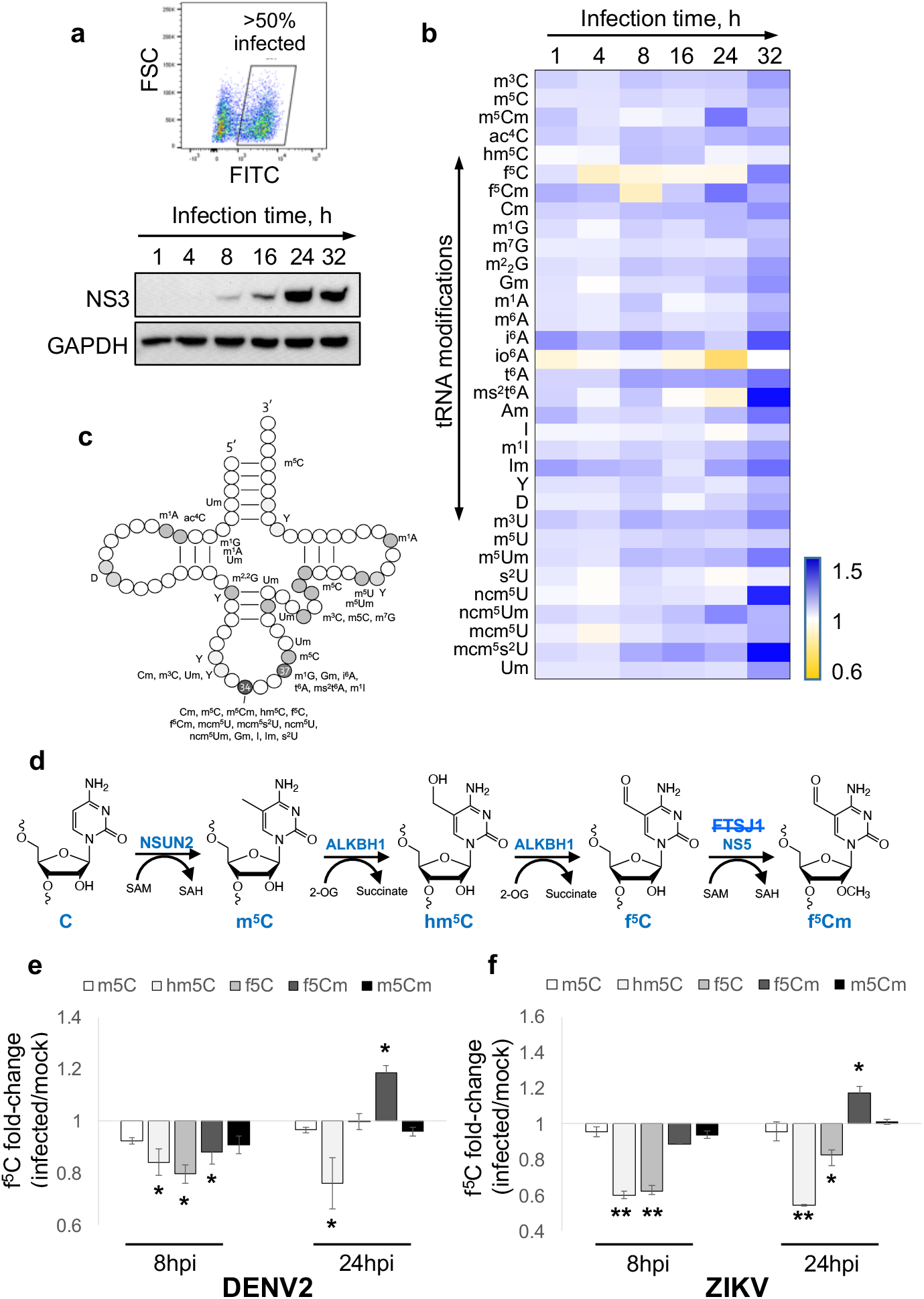
Host tRNA epitranscriptome reprogramming during Flavirus infection. (**a**) Fluorescence-activated cell sorting (FACS) analysis of infected cells using FITC 4G2 staining (upper panel). DENV2 NS3 immunoblot of cell lysates at indicated time points post-DENV2 infection. GAPDH housekeeping protein used as loading control (lower panel). (**b**) Heat map of tRNA modification (rows) changes of DENV2 NGC-infected cells versus mock-infected cells at indicated time points (columns). Gradient scale bar illustrating increased (blue) and decreased (yellow) modification levels as fold-changes in infected versus mock. (**c**) tRNA structural location of the 30 modifications quantified in panel **b**. Some locations are approximate due to varying tRNA structures. (**d**) Enzymes and tRNA modifications involved in f^5^Cm biogenesis: m^5^C, hm^5^C, f^5^C, f^5^Cm, m^5^Cm. NS5 is a DENV-encoded 2’-*O-*methyltransferase. FTSJ1 is the Huh-7 cell 2’-*O*-methyltransferase proposed for f^5^Cm modification in other studies, but not found to participate here (strikethrough). (**e, f**) Changes in levels of f^5^Cm-related tRNA modifications in early (8 hpi) and late (24 hpi) infections by DENV2 EDEN2 (**e**) and ZIKV Brazil strain (**f**). Statistical significance was determined using Student’s t test, *p<0.05; **p<0.01.

### DENV infection alters the host cell tRNA epitranscriptome

To study the effect of DENV infection stress on the tRNA epitranscriptome in Huh-7 cells, paired mock-and DENV2-infected cells were sampled at regular intervals following virus infection and 30 tRNA modifications were quantified by liquid chromatography-coupled tandem mass spectrometry (LC-MS/MS; **Table S1**). The heat map in Figure 1b reveals significant infection-dependent changes in the host cell tRNA epitranscriptome during DENV infection. We observed that many of the modifications that were significantly altered during DENV infection are located in the tRNA anticodon loop and most function in anticodon-codon recognition for proper decoding of mRNAs (Fig. 1c). For example, wobble uridine 5-methoxy-carbonyl-methyl-2-thio (mcm^5^s^2^U) at position 34 of tRNAs for lysine, glutamine, and glutamate was significantly increased (>1.5-fold) during the course of DENV infection, while 5-carbamoylmethyluridine (ncm^5^U) levels peaked (>1.5-fold) at 32 hpi. Similarly, the position 37 modification 2-methylthio-N^6^-threonylcarbamoyladenosine (ms^2^t^6^A) increased 2-fold at 32 hpi. Interestingly, we observed reductions in 5-formylcytidine (f^5^C) and 5-formyl-2′-*O-*methylcytidine (f^5^Cm) levels in early DENV2 NGC infection (8 hpi), but f^5^Cm increased significantly later in infection (24-32 hpi). A pathway for generation of f^5^C and f^5^Cm from m^5^C and its ALKBH1 oxidation product 5-hydroxymethylcytidine (hm^5^C) is shown in Fig. 1d [40, 41], with an analogous conversion of m^5^Cm to 5-hydroxymethyl-2’-*O*-methylcytidine (hm^5^Cm) and f^5^Cm also possible [42]. The generality of this epitranscriptome reprogramming was evident in similar infection time courses performed with the clinical isolate EDEN2 strain of DENV2 (Fig. 1e) and a Brazil strain of Zika virus (Fig. 1f), a *Flavivirus* relative. These studies revealed a consistent pattern of altered f^5^Cm biogenesis during infection: insignificant changes in m^5^C and decreased levels of hm^5^C and f^5^C at 8 hpi, followed by increased levels of m^5^Cm and f^5^Cm by 24 hpi (Fig. 1e, f). We previously observed signature changes in the levels of tRNA modifications caused by specific stresses, which led us to ask if patterns we observed here were due to infection-induced oxidative stress [43] or endoplasmic reticulum (ER) stress [44]. To this end, we exposed Huh-7 cells to agents that cause oxidative (ciprofloxacin) [45] and ER stresses (dihydroartemisinin) [46] followed by measurement of tRNA modification changes. As shown in **Figure S2**, the tRNA epitranscriptome changes caused by these chemical stressors differ significantly from changes caused by DENV2 infection (Fig. 1b). This raised the possibilities that the DENV2-induced reprogramming of tRNA modifications reflects cellular responses to other stresses of viral infection or that they serve to facilitate viral infection.

### tRNA writer ALKBH1 regulates DENV infection

The observation of infection-induced changes in f^5^C and f^5^Cm suggested the involvement of the multifunctional DNA and RNA writer and eraser ALKBH1. As shown in Figure 1d, ALKBH1 is a 2-oxoglutarate-and Fe^2+^-dependent RNA dioxygenase responsible for the hydroxylation of m^5^C to hm^5^C, possibly m^5^Cm to hm^5^Cm, and for the subsequent oxidation of hm^5^C and hm^5^Cm to f^5^C in f^5^Cm, respectively, at the wobble position 34 of cytoplasmic tRNA^Leu(CAA)^ and mitochondrial tRNA^Met^ [40–42]. Other host cell enzymes involved in these transformations include NSUN2 that converts C to m^5^C and FTSJ1 that performs 2’-*O*-methylation [41, 47]. Consistent with reduced f^5^C/f^5^Cm levels during DENV2 and ZIKV infection, immunoblot analysis showed concomitant reduction of ALKBH1 protein levels in DENV2-infected cells (Fig. 2a), with the presence of NS5 protein verifying viral replication. Our observation points to ALKBH1-mediated reprogramming of f^5^C-modified cellular tRNAs during flavivirus infection.

**Figure 2.**
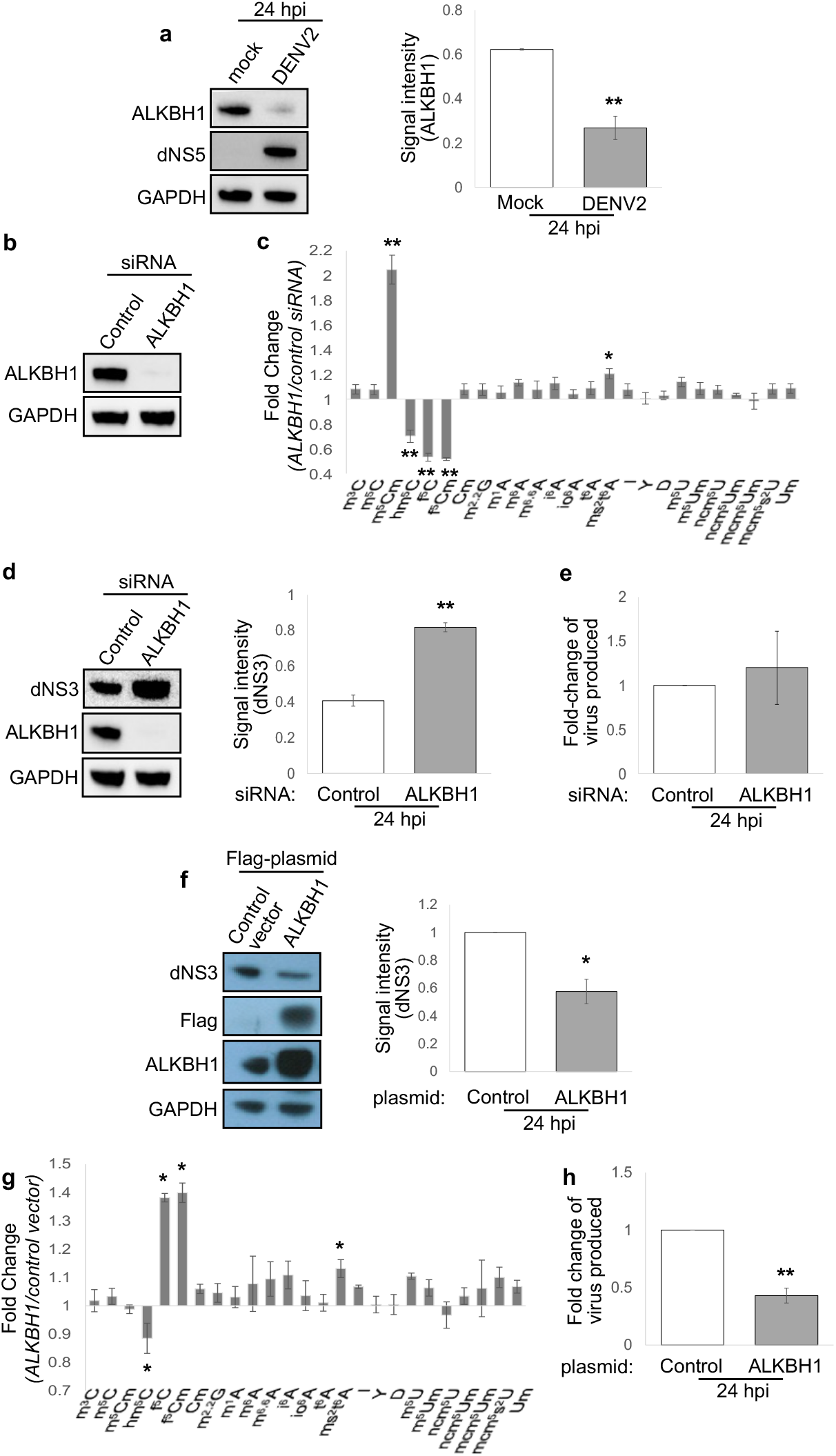
f5Cm writer ALKBH1 restricts DENV2 infection. (**a**) ALKBH1 and DENV2 NS5 immunoblots of DENV2 NGC-and mock-infected cells (left). Densitometric quantitation of NS5 protein (right). (**b**) siRNA knockdown of ALKBH1 assessed by immunoblotting. (**c**) LC-MS/MS profiling of tRNA modifications in ALKBH1 knockdown versus control knockdown cells. (**d**) DENV2 NS3 immunoblot analysis in ALKBH1 knockdown versus control knockdown cells (left). Densitometry quantitation of NS3 protein (right). (**e**) New virus particle production in ALKBH1 knockdown versus control knockdown cells determined by plaque assay. (**f**) DENV2 NS3 immunoblot analysis in ALKBH1-overexpressing versus control cells (left). Densitometry quantitation of NS3 protein (right). (**g**) LC-MS/MS profiling of tRNA modifications in ALKBH1-overexpressing versus control cells. (**h**) New virus particle production at 24 hpi in ALKBH1-overexpressing versus control cells determined by plaque assay. Graphs in all panels show mean ±} SD for 3 biological replicates. All statistical significance was determined using Student’s t test, *p<0.05; **p<0.01.

We next tested the functional role of ALKBH1 in DENV infection by engineering ALKBH1 expression in Huh-7. As shown in Fig. 2b, ALKBH1 was effectively depleted in Huh-7 cells using ALKBH1-specific siRNA, with concomitant reductions of hm^5^C, f^5^C, and f^5^Cm products and increased substrate m^5^Cm (Fig. 2c). As an index of viral replication, dengue NS3 protein levels were increased 24 hpi in the ALKBH1 knockdown cells compared to cells treated with non-targeting control siRNA for both DENV NCG strain (Fig. 2d) and DENV EDEN2 strain (**Fig. S3a**). Quantitation of new virus produced in ALKBH1 knockdown cells indicated a slight but statistically insignificant increase in production of new virus particles over control siRNA-treated cells, suggesting a role for ALKBH1 in modulating viral protein translation though not necessarily leading to new virus production in the cell (Fig. 2e). Conversely, overexpression of ALKBH1 by transient transfection of Huh-7 cells with a FLAG-tagged ALKBH1 plasmid caused increased levels of f^5^C and f^5^Cm products and decreased hm^5^C substrate (Fig. 2g), and also caused significant reductions in both DENV NS3 protein production and new virus particle production (Fig. 2f, h). That compromised fitness of the ALKBH1 engineered cells did not account for these observations was established with cell viability assays (**Fig. S4a, b**). Taken together, these results identify the RNA writer/eraser ALKBH1 as a host restriction factor for DENV replication in human cells, in part by regulating f^5^C/f^5^Cm status of tRNA.

### ALKBH1 and f^5^Cm regulate codon-biased translation of pro-viral transcripts

Given the evidence for ALKBH1’s role in both f^5^Cm biosynthesis in tRNA^Leu(CAA)^ and dengue virus replication, we next sought to define the mechanism linking these activities. Two precedents supported the idea of using proteomics to define this mechanism. First, analogous to the role of ALKBH1-mediated f^5^C in mitochondrial tRNA^Met^ in expanding tRNA decoding from the cognate AUG codon to the non-cognate AUA codon, wobble f^5^Cm in cytoplasmic tRNA^Leu(CAA)^ has been hypothesized to enable decoding of the non-cognate codon UUA in addition to its cognate codon UUG [41]. Second, we previously showed that reprogramming of the tRNA epitranscriptome regulates selective translation of families of codon-biased mRNAs in prokaryotes and eukaryotes [33–38]. We thus assessed the role of ALKBH1 in cellular translation by performing quantitative proteomic analysis of Huh-7 cells with varying levels of ALKBH1 expression and then analyzing codon usage patterns in genes encoding ALKBH1-dependent proteins. For these studies, we created ALKBH1 knockdown cells by transient transfection of ALKBH1-specific siRNA (Fig. 2b**-e**), thus mimicking viral infection, and ALKBH1-complemented cells by introducing an ALKBH1 siRNA-resistant plasmid (pALKr) harboring four consecutive codons that were changed to their synonymous partners at each siRNA-target site in an ALKBH1 knockdown background, thus abolishing siRNA binding (Fig. 3a). This latter cell line controlled for siRNA-mediated cell stress that could affect the epitranscriptome and proteome. Both cell lines showed the expected levels of ALKBH1, though ALKBH1-complemented cells showed approximately 2.5-fold higher ALKBH1 protein expression compared to control Huh-7 cells as determined by quantitative proteomics analysis (Fig. 3b**, Table S2**). Consistent with the role of ALKBH1 in f^5^Cm biosynthesis, tRNA epitranscriptome profiling of ALKBH1-depleted cells showed reduced hm^5^C, f^5^C, and f^5^Cm levels, while ALKBH1-complemented cells showed f^5^C and f^5^Cm levels comparable to that of control siRNA-treated cells (Fig. 3c). Contrary to expectations, however, hm^5^C levels were not restored by ALKBH1 complementation, possibly due to efficient ALKBH1-mediated oxidation of hm^5^C to f^5^C.

**Figure 3.**
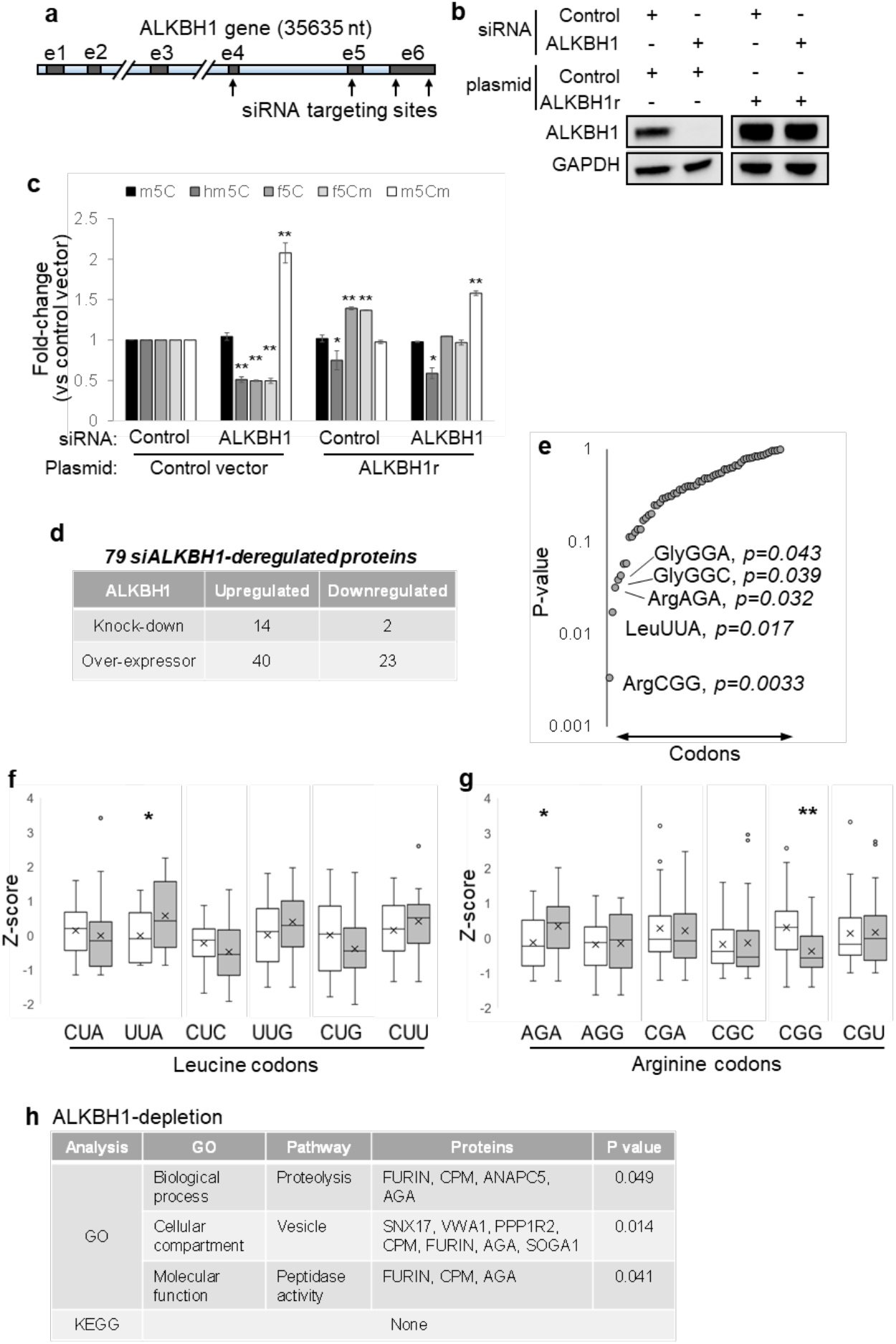
ALKBH1:f5Cm-mediated cellular translational remodeling. (**a**) Illustration of ALKBH1 transcript with indicated siRNA-targeting sites. (**b**) Manipulation of ALKBH1 levels by siRNA knockdown, plasmid overexpression, and complementation using a siRNA-resistant ALKBH1 plasmid as assessed by immunoblotting. (**c**) LC-MS/MS profiling of modifications in f^5^Cm biogenesis of cells with indicated treatments versus cells treated with control siRNA and control plasmid. (**d**) Table of ALKBH1-dependent proteins obtained from proteomics analysis of treated cells from panel **c**. (**e**) Plot of p values derived from analysis of codon Z-scores of ALKBH1 negatively-regulated versus positively-regulated transcripts. (**f,g**) Z-score analysis (number of standard deviations above/below the mean) of leucine (**f**) and arginine (**g**) codons in ALKBH1 negatively-regulated (□) versus ALKBH1 positively-regulated (▪) transcripts. (**h**) Table of GO term and KEGG analysis of proteins deregulated by ALKBH1 depletion assessed by DAVID. Statistical significance was determined using Student’s t test, *p<0.05; **p<0.01.

ALKBH1 knockdown, complemented, and overexpressing cell lines were then subjected to quantitative proteomics analysis by LC-MS with isobaric tags. Here we detected 6315 proteins common to at least two of three biological replicates for all cell lines (**Table S2**). To identify ALKBH1-dependent proteins, we selected proteins that showed a >10% change in expression level in cells treated with ALKBH1 siRNA compared to sham siRNA, and that were further ‘rescued’ by ALKBH1 complementation, where ‘rescue’ was defined by a return of expression levels to baseline, that is, within 5% change in protein expression level in ALKBH1-complemented cells compared to sham siRNA cells. These criteria identified 14 proteins upregulated and 2 proteins downregulated upon ALKBH1 depletion (Fig. 3d). Additionally, we analyzed the ALKBH1 overexpressing cell line for potential ALKBH1-dependent proteins. Here, we similarly selected proteins that showed a >10% change in expression level in cells overexpressing ALKBH1 compared to sham siRNA cells, and further filtered out proteins that showed similar changes in expression in ALKBH1 siRNA cells, that is, >5% change in protein expression level (in the same direction) in ALKBH1 siRNA cells. These criteria identified 40 upregulated proteins and 23 downregulated proteins upon ALKBH1 overexpression (Fig. 3d). Combining both of these analyses yielded a total of 79 proteins that were perturbed by changes in ALKBH1 expression. Of these, 37 proteins were suppressed by ALKBH1 and increased in abundance when ALBKH1 expression decreased, while 42 proteins were induced by ALKBH1, increasing or decreasing directly with ALKBH1 levels (Fig. 3d).

The genes for these 79 ALKBH1-dependent proteins were then assessed for codon usage patterns for 59 codons (excluding STOP codons and Met and Trp that are coded by only one codon) using a gene-specific codon counting algorithm [48]. Strikingly, the f^5^Cm-regulated leucine UUA codon was the second most significantly (p = 0.017) and differentially enriched between the two sets of ALKBH1-dependent proteins, after arginine CGG codon (p = 0.003) (Fig. 3e). Two features of the ALKBH1-dependent proteome were consistent with the previous observation that ALKBH1-mediates f^5^Cm modification of cytoplasmic tRNA^Leu(CAA)^ to expand tRNA decoding from the cognate UUG codon to the non-cognate UUA codon[41] (Fig. 3e, f). First, ALKBH1 depletion enhanced translation of mRNAs lacking the UUA codon, which was replaced by one of the other five leucine codons (UUG, CUU, CUC CUA, CUG) (Fig. 3f). Second, ALKBH1 over-expression enhanced translation of mRNAs enriched with UUA. Surprisingly, the arginine codon CGG was found to be the most significantly (p = 0.003) and differentially enriched between the two sets of ALKBH1 dependent proteins, along with another arginine codon AGA (p = 0.032) (Fig. 3g), although their link to ALKBH1 is unclear at present.

Having identified the unique UUA codon biases in ALKBH1-dependent proteins, we next sought to define the function of these proteins. Here we applied KEGG pathway and GO term analysis to proteins that changed in abundance upon infection-mimicking ALKBH1 depletion. Although no pathways were identified from KEGG analysis, likely due to the small data set analyzed (16 proteins), GO term analysis of proteins deregulated upon ALKBH1 depletion revealed statistically significant enrichment of the terms ‘proteolysis’ (p = 0.049), ‘vesicle’ (p = 0.014) and ‘peptidase activity’ (p = 0.041) in the ‘biological process’, ‘cellular compartment’ and ‘molecular function’ categories, respectively (Fig. 3h). Enrichment of these GO terms in ALKBH1 depletion are consistent with critical steps in viral protein processing (e.g., viral prM cleavage by FURIN) and trafficking during viral replication in cells. This is further consistent with a model in which alteration of ALKBH1 and f^5^Cm leads to increased codon-biased translation of pro-viral transcripts and pro-viral remodeling of the cellular proteome. Importantly, the proteomic analysis of the ALKBH1-knockdown cells, which does not take into account influences of infection, likely reflects only the early stages of infection, when f^5^C and f^5^Cm are reduced and when translation of viral proteins has not reached its peak in dengue-infected cells. At the late stage of 24 hpi, however, f^5^Cm is significantly increased in parallel with dengue virus protein NS5, as discussed next.

### Dengue virus NS5 methylates host cytoplasmic tRNA^Leu(CAA)^ to generate f^5^Cm in late-stage DENV infection

One observation remained unexplained at this point: why are f^5^Cm levels increased at 24 hpi when ALKBH1 expression is suppressed by viral infection? To answer this question, we first defined the wobble modifications present in the tRNA^Leu(CAA)^ isoacceptor in mock-and DENV-infected Huh-7 cells at 24 hpi. Here we used reciprocal circulating chromatography (RCC) to purify the isoacceptor and LC-MS/MS analysis of ribonuclease-digested tRNA to identify and quantify the spectrum of C34 modifications [41]. As shown in Figure 4a, this analysis revealed that, in uninfected cells, hm^5^C (38%), hm^5^Cm (24%), and f^5^Cm (21%) were the major species present in tRNA^Leu(CAA)^, with unmodified C, m^5^C and f^5^C/m^5^Cm accounting for the remaining 17% of C34 modifications, keeping in mind that m^5^C is also present at other locations in some tRNAs. At 24 hpi, however, the relative abundance of hm^5^C decreased by 10%, while f^5^Cm increased by 11%, suggesting an increase in f^5^Cm biosynthesis, possibly due to 2’-*O*-methylation of f^5^C in the virus-infected cells, despite reduced levels of ALKBH1 and f^5^C (Figs. 1b, 4a).

**Figure 4.**
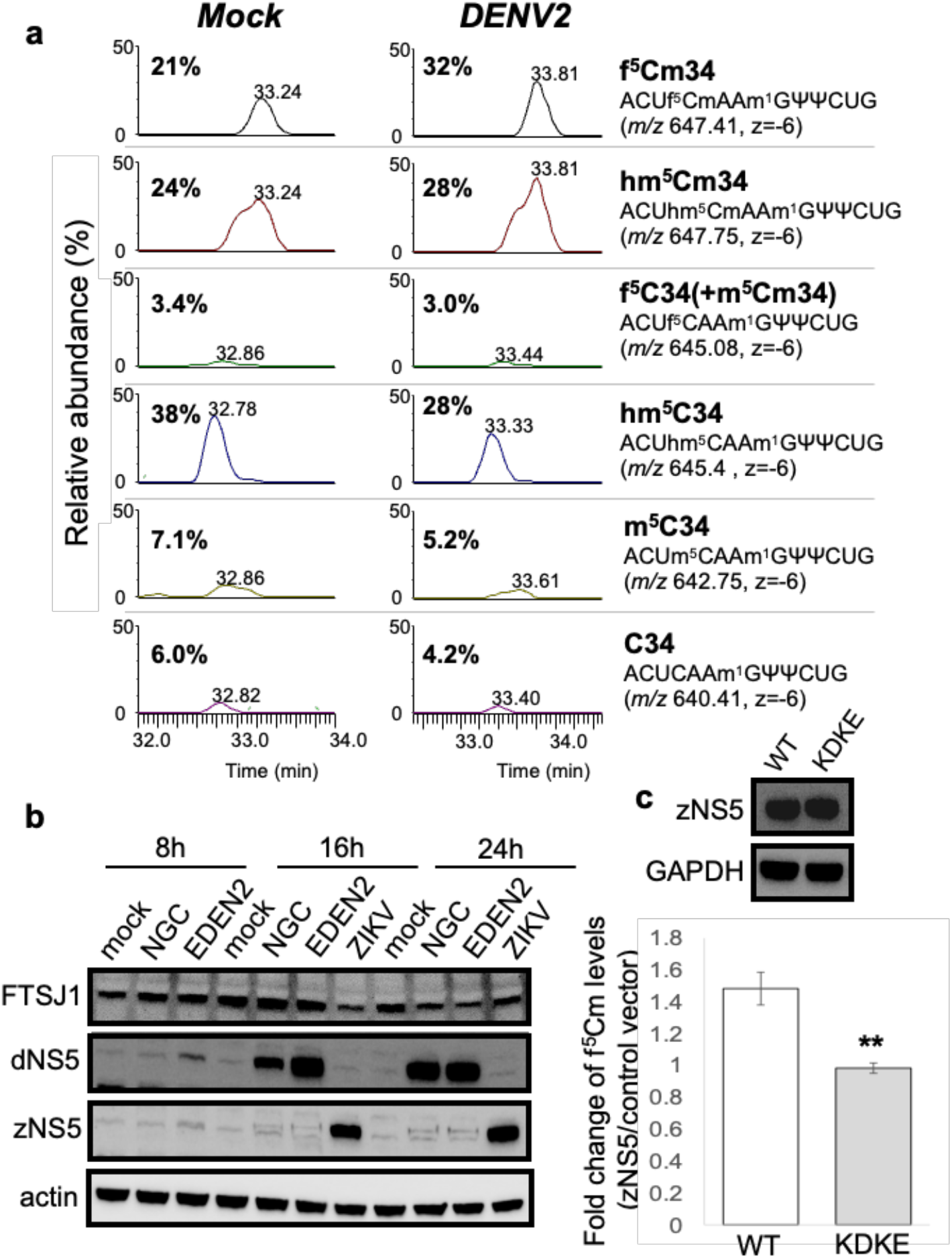
Cytoplasmic tRNA^Leu(CAA)^ is predominantly f^5^Cm-modified in late DENV2 infection. **(a)** Relative abundance of modifications m^5^C, hm^5^C, f^5^C/m^5^Cm, hm^5^Cm and f^5^Cm of isolated ct-tRNA^Leu(CAA)^ from mock-(left panel) and DENV2 NGC-infected Huh-7 cells (middle panel) at 24 hours post-infection. The respective anticodon fragments harboring C34 modifications and *m/z* values are indicated (right panel). **(b)** Human cellular FTSJ1 and viral NS5 immunoblots of cell lysates from indicated time points post-infection of mock-and DENV2 NGC, EDEN2 and ZIKV brazil strain-infected cells. **(c)** ZIKV NS5 (zNS5) immunoblots of wild-type (WT) and Mtase-dead mutant (KDKE) (upper panel). Changes in f^5^Cm modification levels in cells overexpressing WT and KDKE mutant zNS5 compared to control empty vector measured at 24 h post-transfection (lower panel). Statistical significance was determined using Student’s t test, *p<0.05; **p<0.01.

The observation of an infection-induced increase in f^5^Cm in tRNA^Leu(CAA)^ raised the question of which enzyme was responsible for the 2’-*O*-methylation of f^5^C or for 2’-*O*-methylation of m^5^C with subsequent oxidation to f^5^Cm. Human FTSJ1 is a tRNA methyltransferase that is involved in 2’-*O*-methylation of tRNAs at residues 32 and 34, and a study by Kawarada *et al.* demonstrated FTSJ1-mediated 2’-*O*-methylation of cytoplasmic tRNA^Leu(CAA)^ to f^5^Cm/hm^5^Cm [41].

We therefore asked if FTSJ1 was responsible for the increased f^5^Cm-modified cytoplasmic tRNA^Leu(CAA)^ levels found during DENV infection. Proteomics analysis of mock-and DENV-infected Huh-7 cells showed comparable levels of FTSJ1 during early infection (8 hpi) and a slight decrease in FTSJ1 cellular levels by 24 hpi (**Table S3**). Immunoblot analysis of FTSJ1 in mock-, DENV2 NGC-, DENV2 EDEN2-, and ZIKV-infected cells verified the proteomics results, showing comparable FTSJ1 expression at 8 hpi and a reduction in FTSJ1 levels at 24 hpi (Fig. 4b). These results raised the possibility that FTSJ1 was not the primary methyltransferase responsible for the increased f^5^Cm-modified cytoplasmic tRNA^Leu(CAA)^ during DENV infection. Further, since FTSJ1 was reported to methylate both hm^5^C and f^5^C of tRNA^Leu(CAA)^ to hm^5^Cm and f^5^Cm, respectively, but only f^5^Cm-modified cytoplasmic tRNA^Leu(CAA)^ was increased in virus-infected cells, these results suggest the involvement of another methyltransferase in f^5^Cm biogenesis during viral infection.

One candidate for the f^5^Cm-completing 2’-*O*-methyltransferase was the virally-encoded NS5 protein. We had previously shown that this multifunctional enzyme, which installs a host-like m^7^G-and Am-containing 5’-cap on the viral RNA genome and also serves as the replicative polymerase, performs 2’-O-methylation of A, C and G throughout the viral RNA genome and in human rRNAs [49]. To test this idea, we validated NS5 expression by immunoblot analysis of the infection time course for DENV2 strains NGC and EDEN2 and Zika virus, which showed detectable viral protein by ∼16 hpi and maximal protein levels at 24 hpi (Fig. 4b). This time course paralleled f^5^Cm levels at 16 and 24 hpi (Fig. 1b) following DENV2 NGC infection (Fig. 4b). To conclusively establish that viral NS5 was responsible for f^5^Cm biosynthesis during infection, we generated plasmids harboring Zika virus NS5 (wild-type full-length) and the corresponding catalytically-dead mutant in which the tetrad MTase active site residues K61-D146-K181-E217 were substituted with alanine [49]. LC-MS/MS profiling of f^5^Cm levels in purified tRNA from cells transfected with wild-type NS5 showed a 1.5-fold increase in f^5^Cm levels over mock-transfected Huh-7 cells, which was not observed with the catalytically-dead NS5 mutant (Fig. 4c). These results suggest that flavivirus NS5 is capable of contributing to 2’-*O*-methylation of f^5^C to f^5^Cm in human tRNA, thus facilitating codon-biased translation of pro-viral host proteins.

## Discussion

The studies presented here illustrate how a virus not only hijacks the host cell translational machinery to successfully replicate, but also manipulates the host tRNA epitranscriptome. This manipulation of tRNA modifications was detected at two levels. First, the host tRNA writer ALKBH1 functions as a restriction factor for dengue virus infection, with virus-induced silencing of ALKBH1 expression allowing expression of pro-viral host proteins. The second level of epitranscriptome interference occurs with the viral NS5 protein, with this multifunctional protein methylating host tRNA and raising f^5^Cm levels, with yet-to-be defined effects on the proteome of the infected host cell. While there are certainly other factors at play in regulating viral replication in human cells and while we do not know the mechanism by which dengue infection suppresses ALKBH1 expression, we now know that dengue virus exploits the host tRNA epitranscriptome to promote viral replication.

The importance of ALKBH1 as a cellular restriction factor of dengue virus infection in human host cells is underscored by its significant effects on viral protein production and viral replication. Our results suggest that ALKBH1 achieves this regulation by altering part of the cell proteome. While our proteomics analysis of ALKBH1-depleted cells does not account for other infection-induced molecular changes, the observation of ALKBH1-dependent protein expression provides important insights into a subset of molecular changes that may contribute to the viral infection phenotype. For example, reduced ALKBH1 early in the infection caused increased expression of human enzymes known to participate in processing of DENV proteins, such as the endoproteinase FURIN involved in the cleavage of DENV precursor membrane protein (prM) to release a peptide, pr, that facilitates dimerization of E proteins critical to virion assembly [50, 51]. Other host factors associated more generally with viral infections were similarly upregulated by ALKBH1-depletion. These include cellular protein SNX17 that facilitate viral entry and eukaryotic translation initiation factor EIF4H involved in translation of viral transcripts. These results suggest that the virus suppresses ALKBH1 as part of a broader scheme to facilitate viral infection, replication, and release. Furthermore, the fact that ALKBH1 has also been implicated as a demethylase of m^6^A in DNA as well as m^1^A and m^3^C in mRNA of mammalian cells raises the possibility that ALKBH1 is a master stress-response factor that coordinates regulation on multiple epigenetic and epitranscriptomic levels to elicit an appropriate cellular response to virus-induced stress.

So how does viral suppression of ALKBH1 lead to pro-viral changes in the host proteome in dengue infection? The results presented here are consistent with our published model for translational regulation of stress response in which stress-specific epitranscriptomic reprogramming leads to codon-biased translation of response genes needed for adaptation and survival, a mechanism observed in both prokaryotes and eukaryotes [29, 33–36, 39, 52]. Here we presented evidence that ALKBH1-mediated, f^5^Cm-dependent codon-biased translation may contribute to the response of human host cells to dengue virus infection. Specifically, during the early stages of dengue virus infection, reduced cellular levels of ALKBH1 lead to lower levels of wobble f^5^C and f^5^Cm34 in tRNA^Leu(CAA)^. The proteomics data in the ALKBH1 knockdown cell line suggests that reduced ALKBH1 activity in infected cells reduces the capacity of the tRNA pool to read the UUA codon in mRNA, which in turn reduces translation of UUA-enriched transcripts and increases translation of mRNAs lacking UUA and enriched with other synonymous codons for leucine. Similarly, when ALKBH1 is over-expressed, f^5^C and f^5^Cm levels in tRNA^Leu(CAA)^ increase, which leads to enhanced translation of mRNAs enriched with UUA codon. This is consistent with previously observed codon-biased translation regulated by tRNA modifications [29, 34–36, 38, 39, 52] which is now being observed in human cells [29, 32]. While we cannot rule out a role for the ALKBH1-mediated f^5^C axis in mitochondrial tRNA^Met^ in regulating mitochondrial translation, this is unlikely to be the case as analysis of mitochondrial tRNA^Met^ by RCC showed that it was almost completely f^5^C-modified in both mock-and DENV2-infected cells (**Fig. S5**). This points instead towards the ALKBH1:f^5^Cm34 regulation of cytoplasmic tRNA^Leu(CAA)^ in codon-biased translational remodeling that influences DENV replication. Interestingly, UUA codon frequencies were similar between the genomes of DENV and humans (**Fig. S3b**), based on genome averages, but gene specific differences are likely playing important roles in the regulation of specific pathways.

A recent publication by Jungfleisch et al. [53] that focused only on two modifications showed increases in mcm^5^U and decreases in mcm^5^s^2^U with Chikungunya virus (CHIKV) infection of HEK293, which they linked to infection-induced increases in KIAA1456 (hTRM9L or TRMT9B) [53]. The pathways for formation of mcm^5^U and mcm^5^s^2^U are well established by several groups to involve ALKBH8/TRMT112 and CTU1/CTU2, with no involvement of KIAA1456/hTRM9L/TRMT9B, which lacks catalytic activity for these modifications [54–56]. Further, we demonstrated that KIAA1456/hTRM9L/TRMT9B is stress-induced phosphosignaling protein [55, 56], which raises the possibility that changes to mcm^5^U and mcm^5^s^2^U during CHIKV infection are part of a larger coordinated stress response program. The extensive epitranscriptomic reprogramming we observed during dengue virus infection supports the idea that there is a coordinated response to that regulates translation.

It is possible that the ALKBH1-mediated translation and consequent accumulation of viral NS5 during early dengue infection may present a viral protein switch from translation to replication of the viral genome – two mutually exclusive events of the virus life cycle that use the plus strand viral genome. Our findings that ALKBH1-mediated reduction in f^5^C/f^5^Cm34 levels (and presumably enhanced translation of mRNAs lacking UUA such as NS5) were detected at 4 to 8 hpi coincides with the observed replication kinetics of dengue virus plus strand-RNA that exhibits a lag phase of approximately 12 hpi followed by the replication phase [57], suggesting a possible epitrancriptomic link between temporal regulation of dengue virus translation and replication.

The combined results of our studies also suggest there is a series of epitranscriptome-mediated proteome shifts along the time course of dengue virus infection. This is illustrated by observations in the early (8 hpi) and late (24 hpi) stages of infection. Early in the infection, down-regulation of ALKBH1 caused reduced f^5^C and f^5^Cm at the wobble position of tRNA^Leu(CAA)^ (Fig. 1), which was mimicked by ALKBH1 knockdown in Huh-7 cells (Fig. 2c). The latter studies also showed increases in m^5^Cm, which may indicate that m^5^Cm is a substrate for ALKBH1 to form hm^5^Cm and then f^5^Cm (Fig. 1d). These changes in the tRNA^Leu(CAA)^ modification program were associated with a reduction in the translation of genes with UUA codon enrichment, which are enriched with pro-viral pathways (Fig. 3). This is consistent with the observation that ALKBH1 levels regulate dengue virus replication, with reduced ALKBH1 facilitating viral protein production (Fig. 2). There were not many other major changes in tRNA modifications observed at 8 hpi (Fig. 1b), but we only examined 30 of the ∼50 known human RNA modifications, so others could have been affected by the infection. One modification that showed no changes was m^1^A. In addition to f^5^C and f^5^Cm metabolism, ALKBH1 has been proposed to demethylate m^1^A in tRNA and m^3^C in mRNA [58], and both Suzuki and coworkers and He and coworkers have shown ALKBH1-dependent changes in the level of m^1^A in tRNAs in cells [41, 58]. There are several possible explanations for this discrepancy. First, ALKBH1 may not be the only m^1^A demethylase, with cellular m^1^A levels being regulated site-specifically and by the balance of demethylase activity. Second, m^1^A is located at several positions in about 10 different tRNA isoacceptors and isodecoders, so analysis of m^1^A levels in total tRNA may mask subtle site-specific m^1^A changes in individual tRNAs. Third, as we and others have demonstrated conclusively, m^1^A is exceptionally difficult to quantify accurately due to its adventitious conversion to m^6^A by the Dimroth rearrangement that occurs spontaneously at all steps of RNA isolation and LC-MS/MS analysis [59]. It will be important to continue to quantify the epitranscriptome changes for remaining RNA modifications and for different forms of RNA.

At the 24 hpi late stage of infection, the epitranscriptome changes and shift in codon-biased translation are again consistent with precedent and expectation. Even with reduced levels of ALKBH1 at 24 hpi, the level of f^5^Cm increased significantly late in the infection (Fig. 1b). This increase in f^5^Cm is best explained by the observation of maximal NS5 expression at this late stage of infection (Fig. 4a) and the proof that NS5 is responsible for the increased f^5^Cm and not the host cell FTSJ1 enzyme (Fig. 4b). This increase in f^5^Cm also correlates with enhanced translation of mRNAs enriched with UUA codon, as observed in ALKBH1 over-expressing cells with high f^5^Cm levels in tRNA^Leu(CAA)^ (Fig. 3). Interestingly, in addition to f^5^Cm, there was a significant increase in several other 2’-*O*-methylated RNA modifications at 24 hpi: m^5^Cm, Im, Am, and ncm^5^Um (Fig. 1b). While there were no changes in expression of host 2’-*O*-methyltransferase FTSJ1 that could account for the increases in these tRNA modifications, it is likely that they were catalyzed at least in part by the high levels of NS5 protein at 24 hpi (Fig. 3b). We previously demonstrated that NS5 is capable of 2’-O-methylating both the viral RNA genome and host cell RNAs [49].

While there are certainly many other transcriptional and translational factors at play in dengue virus infection, the results of our studies support the model depicted in Figure 5 and clearly establish that (1) infection reduces ALKBH1, (2) ALKBH1 inversely regulates viral replication, (3) ALKBH1 regulates codon-biased translation of pro-viral genes, and (4) viral NS5 functions as a surrogate for ALKBH1. The sum of these observations supports a mechanistic model in which dengue virus hijacks the host tRNA epitranscriptome to promote viral replication.

**Figure 5.**
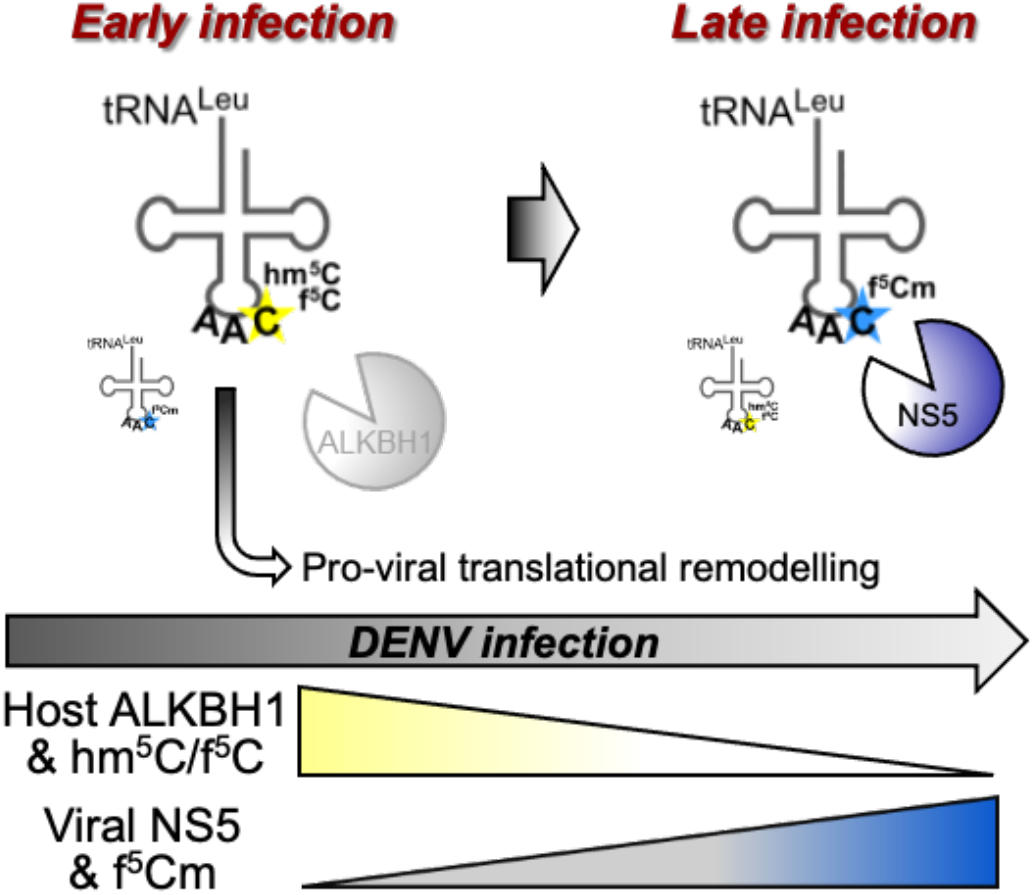
Model of ALKBH1:f^5^Cm-mediated translational remodeling in DENV infection. In early stages of DENV infection, host cellular ALKBH1 levels decrease resulting in reduced f^5^Cm-modified ct-tRNA^Leu(CAA)^ reduced decoding of non-cognate UUA codon. Transcripts lacking UUA are consequently preferentially translated and many of which have virus-associated functions, resulting in pro-viral translational remodeling of the host proteome to facilitate viral replication. As the virus replicates and translates its own mRNA, accumulating the MTase NS5 in the cell at later stages of infection, NS5 contributes to f^5^Cm-modification of tRNA, increasing UUA decoding and further cellular translational remodeling that may contribute to viral exocytosis.

## Methods

### Cells and viruses

Huh-7 and BHK21 cells were cultured in Dulbecco’s Modified Eagle Medium (Thermo Fisher Scientific). C6/36 cells were cultured in Roswell Park Memorial Institute (RPMI) 1640 Medium (Thermo Fisher Scientific). Media were supplemented with 10% fetal bovine serum (Gibco) and cells were cultured in 5% CO2 at 37 °C. Dengue serotype 2 New Guinea C and EDEN2 strains were gifts from Drs. Ooi Eng Eong and Subhash Vasudevan, respectively, Duke-NUS Medical School. Viruses were propagated in C6/36 cells and titrated by plaque assay in BHK21 cells.

### Viral infections

Huh-7 cells were seeded one day prior to infection at 70% confluency. Cells were infected with dengue serotype 2 NGC strain at MOI 1 for 1 h at 37 °C with intermittent shaking. Cells were washed with PBS, grown in fresh medium and harvested at the indicated times.

### Flow cytometry

The proportion of dengue-infected cells was assessed by intracellular staining of dengue E-protein and flow cytometry. Cells were fixed with 4% paraformaldehyde and permeabilized with 0.1% saponin (Sigma-Aldrich). Cells were stained with mouse monoclonal 4G2 antibody (Merck Millipore) and AlexaFluor488 anti-mouse secondary antibody (Thermo Fisher Scientific), and subsequently analyzed on a LSRII flow cytometer (BD Biosciences).

### Total RNA isolation and tRNA purification

Total RNA was obtained from cell pellets using phenol-chloroform extraction and clean up using PureLink miRNA kit (Invitrogen). Integrity of total RNA was assessed by bioanalyzer (Agilent Series 2100). tRNA was purified by size exclusion chromatography, using the Bio SEC-3 300A HPLC column (Agilent) and an isocratic elution with 100 mM ammonium acetate (pH 7.0, 60 °C) on an Agilent 1200 HPLC system.

### tRNA modification detection and quantitation

Purified human tRNA was enzymatically hydrolyzed into ribonucleosides in the presence of deaminase inhibitors and antioxidants as described previously^46,28,47^. Digested ribonucleosides were separated on Hypersil GOLD aQ reverse-phase column (3 µm particle size; 150 x 2.1 mm, Thermo Fisher Scientific) and detected using Agilent 6490 triple quadrupole LC/MS mass spectrometer in positive-ion mode. Relative quantities of each ribonucleoside were assessed as a function of viral infection/transfection treatment.

### tRNA isolation and mapping

Total RNA was extracted from mock-and DENV2-infected Huh-7 cells as described above. ct-tRNA^Leu(CAA)^ and mt-tRNA^Met^ were isolated by reciprocal circulating chromatography (RCC) using 5΄-terminal ethylcarbamate amino-modified DNA probes described previously^30^. Isolated tRNA (1 pmol) was digested with RNase T1 and subjected to capillary liquid chromatography (LC)–nanoelectrospray ionization (ESI)–mass spectrometry (MS) as described^30^.

### Plasmids

Sequential site-directed mutagenesis was performed using wild-type ALKBH1 plasmid (OHu05179, GenScript) as a template with the following primer pairs: site 1, 5’- GTTCCTGAGATACAAGGAGGCAACTAAACGGAGACCCCGAAGTTTAC-3’ and 5’- CTCCGTTTAGTTGCCTCCTTGTATCTCAGGAACTCTTTGCTCTGTTC-3’; site 2, 5’- GTTTATGCACAGCGGCGATATTATGATAATGTCGGGTTTCAGCCGC-3’ and 5’- CGACATTATCATAATATCGCCGCTGTGCATAAACATGGCCGTGGGGGC-3’; site 3, 5’- CAGCTACTTGAAAACAGCCCGCGTTAACATGACTGTCCGACAGGTC-3’ and 5’- CAGTCATGTTAACGCGGGCTGTTTTCAAGTAGCTGGCACACACCTGC-3’; site 4, 5’- CTGCCATCTGGACGATCAAAACAGCGAAGTAAAACGGGCCAGGATAAAC-3’ and 5’- CGTTTTACTTCGCT<u>G</u>TT<u>T</u>TG<u>A</u>TC<u>G</u>TCCAGATGGCAGAAACCTTCTGTAC-3’, where synonymous mutations of the four consecutive codons at each siRNA-targeting site are underlined. High fidelity Pfu polymerase extension was performed using the following parameters: 94 °C for 30 s, followed by 17 cycles of 94 °C for 30 s, 55 °C for 1 min, 68 °C for 7 min with subsequent DpnI digestion for 1 h at 37 °C. The digested products were transformed, screened by colony PCR and introduced mutations were confirmed by sequencing. Similarly, Zika NS5 MTase catalytic-dead KDKE mutant was generated by sequential site-directed mutagenesis using isolated zika virus Brazil strain genome as template with the following primer pairs: K61A: 5’- CGAGGCTCAGCA<u>GC</u>ACTGAGATGGTTCGTCGAGAGAAATATGGTC-3’ and 5’- GAACCATCTCAGTGCTGCTGAGCCTCGCGACACAGCGTGATGGTC-3’; D146A: 5’- CATTGTTGTGTGCAATAGGGGAGTCGTCACCAAATCCCAC-3’ and 5’- GACTCCCCTATTGCACACAACAATGTGTCACACTTTTCTGGC-3’; K182A: 5’- CCCAATTTTGCATAGCAGTTCTCAACCCATACATGCCCTCAGTC-3’ and 5’- GTATGGGTTGAGAACTGCTATGCAAAATTGGGTGTTGTTGTTCAACC-3’; E217: 5’- ACTCCACACATGCAATGTACTGGGTATCCAATGCCTCCGGGAAC-3’ and 5’- CCCAGTACAT<u>TG</u>CATGTGTGGAGTTTCGTGAGAGTGGATTCCTC-3’, where mutations introduced are underlined.

### Transfections

Huh-7 cells were seeded one day prior to siRNA and plasmid transfections at 70% and 90% confluency respectively. Transfections were performed using 25 nM ALKBH1-specific (8846, Dharmacon) or non-targeting control (D-001810-10-05, Dharmacon) siRNA, 500 ng plasmid DNA and lipofectamine 3000 (Invitrogen) as indicated, according to the manufacturer’s instructions. Knock-down or over-expression of the respective proteins were confirmed by western blot analysis.

### Immunoblotting

Cells were lysed and cell lysates clarified by centrifugation and quantified by Bradford assay. Each cell lysate sample (10 µg of protein) was denatured by boiling in loading buffer for 10 min and subjected to SDS-polyacrylamide gel electrophoresis (SDS-PAGE). Western blot analysis was performed using primary antibodies rabbit anti-ALKBH1 (ab126596, Abcam, Cambridge, UK, 1:1000), mouse anti-GAPDH (sc-47724, Santa Cruz Biotechnology, 1:10000), rabbit anti-dengue NS3 (PA532199, Thermo 1:1000), rabbit anti-dengue NS5 (GTX103350, GeneTex, 1:1000), rabbit anti-zika NS5 (GTX133312, GeneTex, 1:1,000), anti-FTSJ1 (11620-1-AP, Proteintech, 1:1000) and secondary antibodies peroxidase conjugated goat anti-rabbit (A6154, Sigma-Aldrich, 1:10000) and goat anti-mouse (Sigma-Aldrich, 1:10000) in 2% milk.

### Proteomics

All sample preparation and proteomics analysis were performed by the Proteomics Core Facility at Nanyang Technological University, Singapore. To prepare proteins in solution, 100 μg of protein from each condition was subjected to in-solution digestion before labelling the resultant peptides using the TMT-10plex Isobaric Label Reagent Set (Thermo Fisher Scientific) according to the manufacturer’s protocol. The labeled samples were combined prior to fractionation on a Xbridge C18 column (4.6 × 250 mm, Waters) and subsequent analysis by chromatography-coupled Orbitrap mass spectrometry.

Fractionated peptides were separated and analyzed using a Dionex Ultimate 3000 RSLCnano system coupled to an Oribtrap Q Exactive mass spectrometer (Thermo Fisher Scientific). Separation was performed on a Dionex EASY-Spray 75 μm × 10 cm column packed with PepMap C18 3 μm, 100 Å (Thermo Fisher Scientific) using solvent A (0.1% formic acid) and solvent B (0.1% formic acid in 100% ACN) at flow rate of 300 nL/min with a 60 min gradient. Peptides were analyzed on a Q Exactive apparatus with an EASY nanospray source (Thermo Fisher Scientific) at an electrospray potential of 1.5 kV. A full MS scan (350–1,600 m/z range) was acquired at a resolution of 70,000 and a maximum ion accumulation time of 100 ms.

Dynamic exclusion was set as 30 s. The resolution of the higher energy collisional dissociation (HCD) spectra was set to 350,00. The automatic gain control (AGC) settings of the full MS scan and the MS2 scan were 5E6 and 2E5, respectively. The 10 most intense ions above the 2,000-count threshold were selected for fragmentation in HCD, with a maximum ion accumulation time of 120 ms. An isolation width of 2 *m/z* was used for MS2. Single and unassigned charged ions were excluded from MS/MS. For HCD, the normalized collision energy was set to 30%. The underfill ratio was defined as 0.3%. Raw data files from the three technical replicates were processed and searched using Proteome Discoverer 2.1 (Thermo Fisher Scientific). The raw LC-MS/MS data files were loaded into Spectrum Files (default parameters set in Spectrum Selector) and TMT 10-plex was selected for the Reporter Ion Quantifier. The SEQUEST algorithm was then used for data searching to identify proteins using the following parameters; missed cleavage of two; dynamic modifications were oxidation (+15.995 Da) (M). The static modifications were TMT-10plex (+229.163 Da) (any N-terminus and K) and Carbamidomethyl (+57 Da) (C). The false discovery rate for protein identification was <1%. The Normalization mode was set based on total peptide amount.

### Statistical analysis

All experiments were performed in with a minimum of two technical replicates and 3 biological replicates. Statistical significance was determined using Student’s t test, *p<0.05; **p<0.01.

## Data Availability

Proteomics data are available at the ProteomeXchange with identifier PXD028029 (http://www.ebi.ac.uk/pride). Mass spectrometry data are included as **Supporting Data Table S1**.

## Acknowledgements

The authors acknowledge funding by National Research Foundation of Singapore through the Singapore-MIT Alliance for Research and Technology Antimicrobial Resistance IRG (PCD) and by grants from the National Institutes of Health (R01 ES026856 and R01 ES024615 to TJB).

## Author contributions

CC, PCD, and TJB conceived of the project, designed the studies, and interpreted data. NSKS performed proteomics analyses. OT, YS, and TS performed tRNA modification mapping. All authors participated in the writing of the manuscript.

## Competing interests

The authors declare no competing interests.

## Supporting Information

**Fig. S1.**
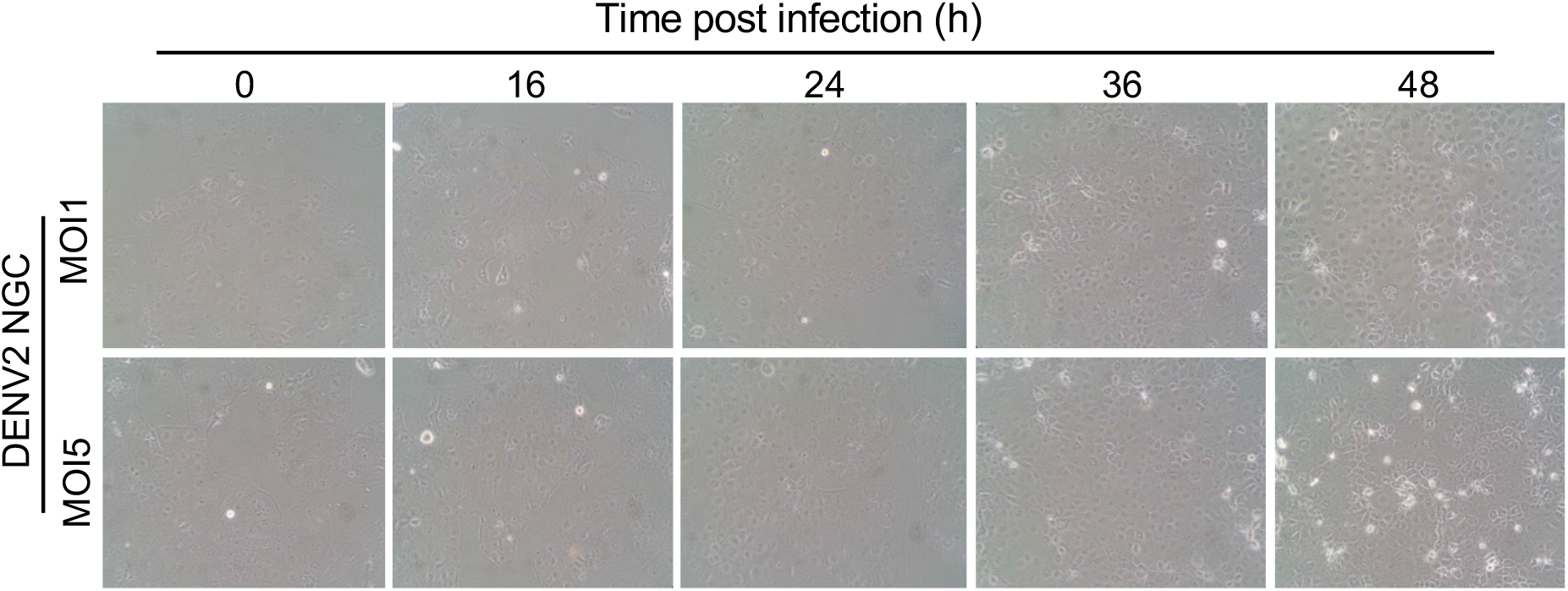
Phase contrast images (10x magnification) of cells infected with DENV2 NGC at MOI 1 and 5, at indicated times post-infection.

**Fig. S2.**
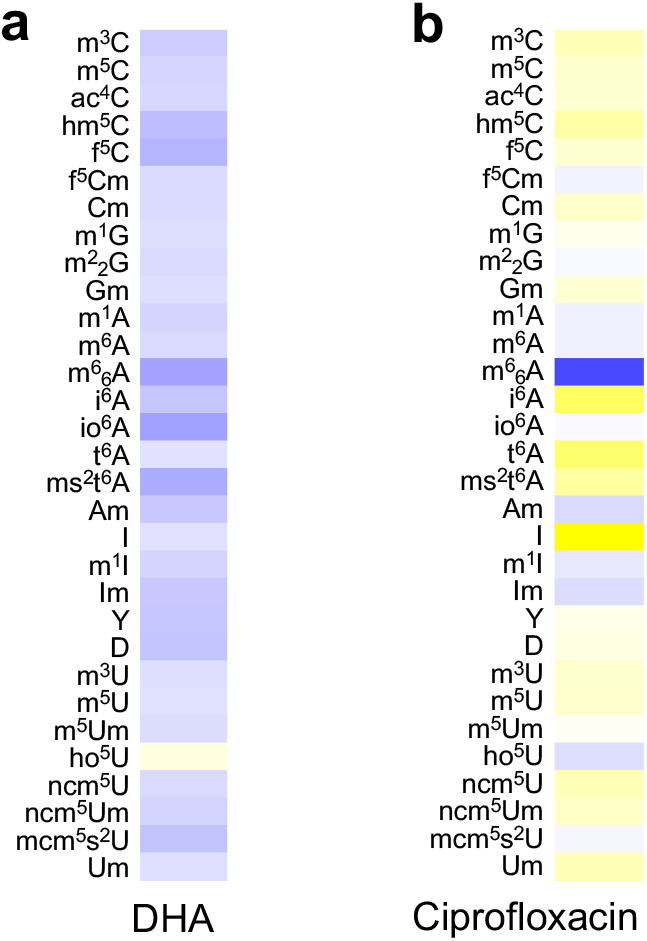
tRNA modification profiles of Huh-7 cells at 24 h following treatment with (**a**) 10 µM dihydroartemisinin (DHA) and (**b**) 20 µM ciprofloxacin assessed by LC-MS/MS.

**Fig. S3.**
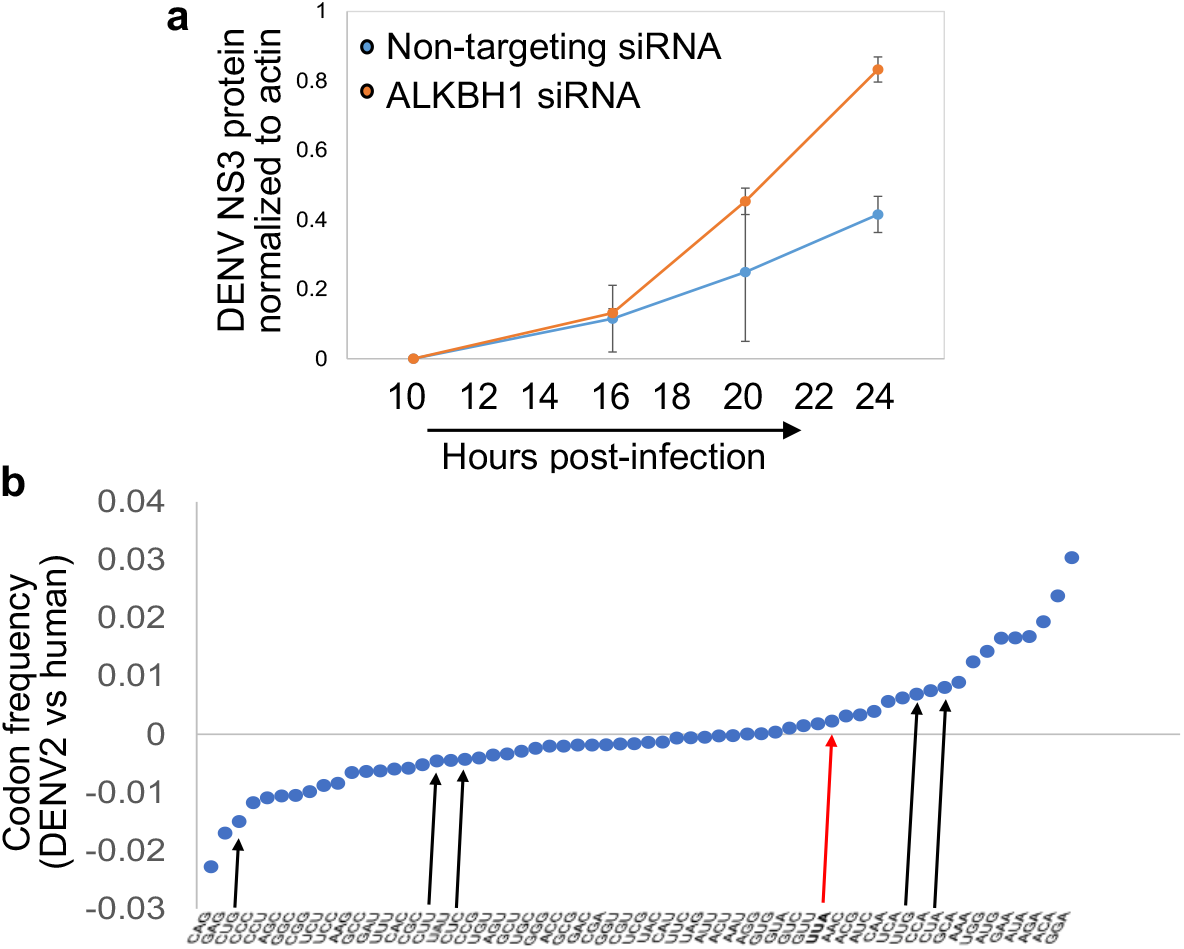
**(a)** Densitometric quantitation of NS3 protein levels at indicated times after infection of Huh-7 cells with DENV strain EDEN2 infection. Huh-7 cells were pre-treated with control non-targeting siRNA (blue circles) and ALKBH1-specific siRNA (orange circles). **(b)** Differences in codon usage frequencies in DENV2 NGC strain versus human host genome. Leucine UUA codon (red arrow) is nearly equally represented in both human and DENV2 genomes, while other leucine codons (black arrows) are not shared equally.

**Fig. S4.**
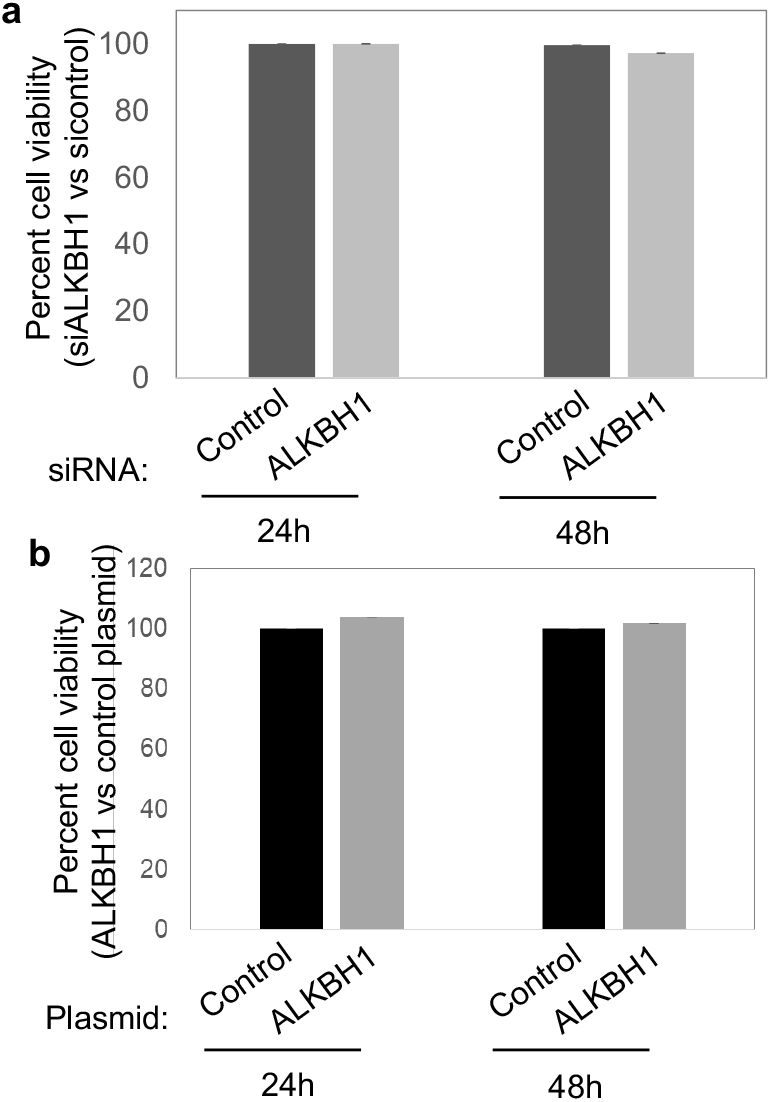
Cell viability determined by MTT assay of Huh-7 cells treated with **(a)** non-targeting and ALKBH1-specific siRNA, and **(b)** control empty vector and ALKBH1 plasmid, at 24h and 48h post-transfection.

**Fig. S5.**
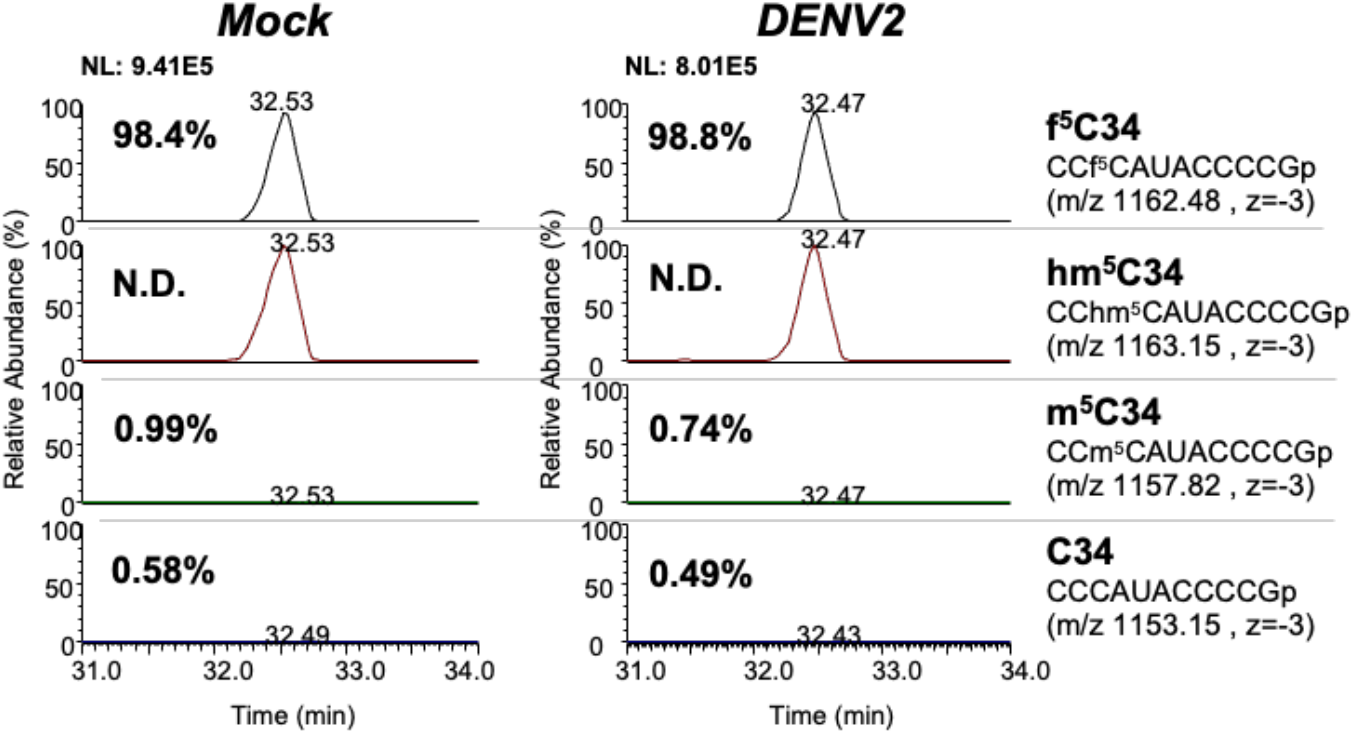
Relative abundance of modifications m^5^C, hm^5^C and f^5^C of isolated mitochondrial tRNA^Met^ from mock-(left panel) and DENV2 NGC-infected Huh-7 cells (middle panel). The respective anticodon fragments harboring C34 modifications and m/z values are indicated (right panel). N.D., not detected.

**Supporting Table S1**: LC-MS analysis of RNA modifications; separate spreadsheet

**Supporting Data Table S2**: Proteomics data for ALKBH1 knockdown and over-expression versus control; separate spreadsheet

**Supporting Data Table S3**: Proteomics data for DENV infection at 8 and 24 hpi; separate spreadsheet

## Notes

### Competing Interest Statement

The authors have declared no competing interest.

## References

1. Mesev EV, LeDesma RA, Ploss A. Decoding type I and III interferon signalling during viral infection. Nat Microbiol. 2019;4(6):914–24.

2. Barrat FJ, Crow MK, Ivashkiv LB. Interferon target-gene expression and epigenomic signatures in health and disease. Nat Immunol. 2019;20(12):1574–83.

3. Wolf D, Goff SP. Host restriction factors blocking retroviral replication. Annu Rev Genet. 2008;42:143–63.

4. Zhou LY, Zhang LL. Host restriction factors for hepatitis C virus. World J Gastroenterol. 2016;22(4):1477–86.

5. Bogerd HP, Doehle BP, Wiegand HL, Cullen BR. A single amino acid difference in the host APOBEC3G protein controls the primate species specificity of HIV type 1 virion infectivity factor. Proc Natl Acad Sci U S A. 2004;101(11):3770–4.

6. Bogerd HP, Wiegand HL, Doehle BP, Lueders KK, Cullen BR. APOBEC3A and APOBEC3B are potent inhibitors of LTR-retrotransposon function in human cells. Nucleic Acids Res. 2006;34(1):89–95.

7. Sayah DM, Sokolskaja E, Berthoux L, Luban J. Cyclophilin A retrotransposition into TRIM5 explains owl monkey resistance to HIV-1. Nature. 2004;430(6999):569-73.

8. Neufeldt CJ, Cortese M, Acosta EG, Bartenschlager R. Rewiring cellular networks by members of the Flaviviridae family. 2018; 3:125–142.

9. Ravindran MS, Bagchi P, Cunningham CN, Tsai B. Opportunistic intruders: how viruses orchestrate ER functions to infect cells. Nat Rev Microbiol. 2016;14(7):407–20.

10. Acosta EG, Bartenschlager R. The quest for host targets to combat dengue virus infections. Curr Opin Virol. 2016;20:47–54.

11. Bhatt S, Gething PW, Brady OJ, Messina JP, Farlow AW, Moyes CL, et al. The global distribution and burden of dengue. Nature. 2013;496(7446):504-7.

12. Guzman MG, Halstead SB, Artsob H, Buchy P, Farrar J, Gubler DJ, et al. Dengue: a continuing global threat. Nat Rev Microbiol. 2010;8(12 Suppl):S7-16.

13. Brai A, Fazi R, Tintori C, Zamperini C, Bugli F, Sanguinetti M, et al. Human DDX3 protein is a valuable target to develop broad spectrum antiviral agents. Proc Natl Acad Sci U S A. 2016;113(19):5388–93.

14. Chen Y, Maguire T, Hileman RE, Fromm JR, Esko JD, Linhardt RJ, et al. Dengue virus infectivity depends on envelope protein binding to target cell heparan sulfate. Nat Med. 1997;3(8):866–71.

15. Heaton NS, Perera R, Berger KL, Khadka S, Lacount DJ, Kuhn RJ, et al. Dengue virus nonstructural protein 3 redistributes fatty acid synthase to sites of viral replication and increases cellular fatty acid synthesis. Proc Natl Acad Sci U S A. 2010;107(40):17345–50.

16. Osuna-Ramos JF, Reyes-Ruiz JM, Del Angel RM. The Role of Host Cholesterol During Flavivirus Infection. Front Cell Infect Microbiol. 2018;8:388..00388. PubMed PMID: 30450339; PubMed Central PMCID: PMCPMC6224431.

17. Tassaneetrithep B, Burgess TH, Granelli-Piperno A, Trumpfheller C, Finke J, Sun W, et al. DC-SIGN (CD209) mediates dengue virus infection of human dendritic cells. J Exp Med. 2003;197(7):823–9.

18. Saiz JC, Oya NJ, Blazquez AB, Escribano-Romero E, Martin-Acebes MA. Host-Directed Antivirals: A Realistic Alternative to Fight Zika Virus. Viruses. 2018;10(9).

19. Zakaria MK, Carletti T, Marcello A. Cellular Targets for the Treatment of Flavivirus Infections. Front Cell Infect Microbiol. 2018;8:398.

20. Suzuki T. The expanding world of tRNA modifications and their disease relevance. Nat Rev Mol Cell Biol. 2021;22(6):375–92.

21. Brocard M, Ruggieri A, Locker N. m6A RNA methylation, a new hallmark in virus-host interactions. J Gen Virol. 2017;98(9):2207–14.

22. Dang W, Xie Y, Cao P, Xin S, Wang J, Li S, et al. N(6)-Methyladenosine and Viral Infection. Front Microbiol. 2019;10:417.

23. Williams GD, Gokhale NS, Horner SM. Regulation of Viral Infection by the RNA Modification N6-Methyladenosine. Annu Rev Virol. 2019;6(1):235–53.

24. Lichinchi G, Zhao BS, Wu Y, Lu Z, Qin Y, He C, et al. Dynamics of Human and Viral RNA Methylation during Zika Virus Infection. Cell Host Microbe. 2016;20(5):666–73.

25. Gokhale NS, McIntyre ABR, McFadden MJ, Roder AE, Kennedy EM, Gandara JA, et al. N6-Methyladenosine in Flaviviridae Viral RNA Genomes Regulates Infection. Cell Host Microbe. 2016;20(5):654–65.

26. Chan C, Pham P, Dedon PC, Begley TJ. Lifestyle modifications: coordinating the tRNA epitranscriptome with codon bias to adapt translation during stress responses. Genome Biol. 2018;19(1):228.

27. Jonkhout N, Tran J, Smith MA, Schonrock N, Mattick JS, Novoa EM. The RNA modification landscape in human disease. RNA. 2017;23(12):1754–69.

28. Chen X, Li A, Sun BF, Yang Y, Han YN, Yuan X, et al. 5-methylcytosine promotes pathogenesis of bladder cancer through stabilizing mRNAs. Nat Cell Biol. 2019;21(8):978–90.

29. Rapino F, Delaunay S, Rambow F, Zhou Z, Tharun L, De Tullio P, et al. Codon-specific translation reprogramming promotes resistance to targeted therapy. Nature. 2018;558(7711):605-9.

30. The Cancer Genome Atlas. 2021. Available from: https://www.cancer.gov/tcga.

31. Zhang Z, Ruan H, Liu CJ, Ye Y, Gong J, Diao L, et al. tRic: a user-friendly data portal to explore the expression landscape of tRNAs in human cancers. RNA Biol. 2020;17(11):1674–9.

32. Orellana EA, Liu Q, Yankova E, Pirouz M, De Braekeleer E, Zhang W, et al. METTL1-mediated m(7)G modification of Arg-TCT tRNA drives oncogenic transformation. Mol Cell. 2021;81(16):3323–38 e14.

33. Begley U, Dyavaiah M, Patil A, Rooney JP, DiRenzo D, Young CM, et al. Trm9-catalyzed tRNA modifications link translation to the DNA damage response. Mol Cell. 2007;28(5):860–70.

34. Chan CT, Deng W, Li F, DeMott MS, Babu IR, Begley TJ, et al. Highly Predictive Reprogramming of tRNA Modifications Is Linked to Selective Expression of Codon-Biased Genes. Chem Res Toxicol. 2015;28(5):978–88.

35. Chan CT, Pang YL, Deng W, Babu IR, Dyavaiah M, Begley TJ, et al. Reprogramming of tRNA modifications controls the oxidative stress response by codon-biased translation of proteins. Nat Commun. 2012;3:937.

36. Chionh YH, McBee M, Babu IR, Hia F, Lin W, Zhao W, et al. tRNA-mediated codon-biased translation in mycobacterial hypoxic persistence. Nat Commun. 2016;7:13302.

37. Deng W, Babu IR, Su D, Yin S, Begley TJ, Dedon PC. Trm9-Catalyzed tRNA Modifications Regulate Global Protein Expression by Codon-Biased Translation. PLoS Genet. 2015;11(12):e1005706.

38. Endres L, Begley U, Clark R, Gu C, Dziergowska A, Malkiewicz A, et al. Alkbh8 Regulates Selenocysteine-Protein Expression to Protect against Reactive Oxygen Species Damage. PLoS One. 2015;10(7):e0131335.

39. Fernandez-Vazquez J, Vargas-Perez I, Sanso M, Buhne K, Carmona M, Paulo E, et al. Modification of tRNA(Lys) UUU by elongator is essential for efficient translation of stress mRNAs. PLoS Genet. 2013;9(7):e1003647.

40. Haag S, Sloan KE, Ranjan N, Warda AS, Kretschmer J, Blessing C, et al. NSUN3 and ABH1 modify the wobble position of mt-tRNAMet to expand codon recognition in mitochondrial translation. EMBO J. 2016;35(19):2104–19.

41. Kawarada L, Suzuki T, Ohira T, Hirata S, Miyauchi K, Suzuki T. ALKBH1 is an RNA dioxygenase responsible for cytoplasmic and mitochondrial tRNA modifications. Nucleic Acids Res. 2017;45(12):7401–15.

42. Huber SM, van Delft P, Tanpure A, Miska EA, Balasubramanian S. 2’-O-Methyl-5-hydroxymethylcytidine: A Second Oxidative Derivative of 5-Methylcytidine in RNA. J Am Chem Soc. 2017;139(5):1766–9.

43. Ivanov AV, Bartosch B, Isaguliants MG. Oxidative Stress in Infection and Consequent Disease. Oxid Med Cell Longev. 2017;2017:3496043.

44. Zhang L, Wang A. Virus-induced ER stress and the unfolded protein response. Front Plant Sci. 2012;3:293.

45. Kalghatgi S, Spina CS, Costello JC, Liesa M, Morones-Ramirez JR, Slomovic S, et al. Bactericidal antibiotics induce mitochondrial dysfunction and oxidative damage in Mammalian cells. Sci Transl Med. 2013;5(192):192ra85.

46. Elhassanny AEM, Soliman E, Marie M, McGuire P, Gul W, ElSohly M, et al. Heme-Dependent ER Stress Apoptosis: A Mechanism for the Selective Toxicity of the Dihydroartemisinin, NSC735847, in Colorectal Cancer Cells. Front Oncol. 2020;10:965.

47. Brzezicha B, Schmidt M, Makalowska I, Jarmolowski A, Pienkowska J, Szweykowska-Kulinska Z. Identification of human tRNA:m5C methyltransferase catalysing intron-dependent m5C formation in the first position of the anticodon of the pre-tRNA Leu (CAA). Nucleic Acids Res. 2006;34(20):6034–43.

48. Doyle F, Leonardi A, Endres L, Tenenbaum SA, Dedon PC, Begley TJ. Gene-and genome-based analysis of significant codon patterns in yeast, rat and mice genomes with the CUT Codon UTilization tool. Methods. 2016;107:98–109.

49. Dong H, Chang DC, Hua MH, Lim SP, Chionh YH, Hia F, et al. 2’-O methylation of internal adenosine by flavivirus NS5 methyltransferase. PLoS Pathog. 2012;8(4):e1002642.

50. Stadler K, Allison SL, Schalich J, Heinz FX. Proteolytic activation of tick-borne encephalitis virus by furin. J Virol. 1997;71(11):8475–81.

51. Li L, Lok SM, Yu IM, Zhang Y, Kuhn RJ, Chen J, et al. The flavivirus precursor membrane-envelope protein complex: structure and maturation. Science. 2008;319(5871):1830-4.

52. Bento-Abreu A, Jager G, Swinnen B, Rue L, Hendrickx S, Jones A, et al. Elongator subunit 3 (ELP3) modifies ALS through tRNA modification. Hum Mol Genet. 2018;27(7):1276–89.

53. Jungfleisch J, Bottcher R, Tallo-Parra M, Perez-Vilaro G, Merits A, Novoa EM, et al. CHIKV infection reprograms codon optimality to favor viral RNA translation by altering the tRNA epitranscriptome. Nat Commun. 2022;13(1):4725.

54. Songe-Moller L, van den Born E, Leihne V, Vagbo CB, Kristoffersen T, Krokan HE, et al. Mammalian ALKBH8 possesses tRNA methyltransferase activity required for the biogenesis of multiple wobble uridine modifications implicated in translational decoding. Mol Cell Biol. 2010;30(7):1814–27. 10.1128/MCB.01602-09. PubMed PMID: 20123966; PubMed Central PMCID: PMC2838068.

55. Begley U, Sosa MS, Avivar-Valderas A, Patil A, Endres L, Estrada Y, et al. A human tRNA methyltransferase 9-like protein prevents tumour growth by regulating LIN9 and HIF1-alpha. EMBO Mol Med. 2013;5(3):366–83.

56. Gu C, Ramos J, Begley U, Dedon PC, Fu D, Begley TJ. Phosphorylation of human TRM9L integrates multiple stress-signaling pathways for tumor growth suppression. Sci Adv. 2018;4(7):eaas9184.

57. Reid DW, Campos RK, Child JR, Zheng T, Chan KWK, Bradrick SS, et al. Dengue Virus Selectively Annexes Endoplasmic Reticulum-Associated Translation Machinery as a Strategy for Co-opting Host Cell Protein Synthesis. J Virol. 2018;92(7).

58. Liu F, Clark W, Luo G, Wang X, Fu Y, Wei J, et al. ALKBH1-Mediated tRNA Demethylation Regulates Translation. Cell. 2016;167(3):816–28 e16.

59. Wang J, Alvin Chew BL, Lai Y, Dong H, Xu L, Balamkundu S, et al. Quantifying the RNA cap epitranscriptome reveals novel caps in cellular and viral RNA. Nucleic Acids Res. 2019;47(20):e130.

